# Integrating *N*-glycan and CODEX imaging reveal cell-specific protein glycosylation in healthy human lung

**DOI:** 10.1101/2024.10.08.617274

**Authors:** Dušan Veličković, Jeffrey Purkerson, Harsh Bhotika, Heidie Huyck, Geremy Clair, Gloria S. Pryhuber, Christopher Anderton

## Abstract

*N*-linked glycosylation, the major post-translational modification of cellular proteins, is important for proper lung functioning, serving to fold, traffic, and stabilize protein structures and to mediate various cell-cell recognition events. Identifying cell-specific *N*-glycan structures in human lungs is critical for understanding the chemistry and mechanisms that guide cell-cell and cell-matrix interactions and determining nuanced functions of specific *N*-glycosylation. Our study, which used matrix-assisted laser desorption/ionization (MALDI) mass spectrometry imaging (MSI) combined with co-detection by indexing (CODEX) to reveal the cellular origin of *N*-glycans, is a significant step in this direction. This innovative technological combination enabled us to detect and differentiate *N*-glycans located in the vicinity of cells surrounding airways and blood vessels, parenchyma, submucosal glands, cartilage, and smooth muscles. The potential impact of our findings on future research is immense. For instance, our algorithm for grouping *N*-glycans based on their functional chemical features, combined with identifying group niches, paves the way for targeted studies. We found that fucosylated *N*-glycans are dominant around immune cells, tetra antennary *N*-glycans in the cartilage, high-mannose *N*-glycans surrounding the bronchus originate from associated collagenous structures, complex fucosylated-tetra antennary-polylactosamine *N*-glycans are spread over smooth muscle structures and in epithelial cells surrounding arteries, and *N*-glycans with Hex:6 HexNAc:6 compositions, which, according to our algorithm, can be ascribed to either tetra antennary or bisecting *N*-glycan, are highly abundant in the parenchyma. The findings suggest cell or region-specific functions for these localized glycan structures.

## INTRODUCTION

Protein *N*-glycosylation is a ubiquitous post-translational modification that plays essential biological roles in vital molecular processes ranging from protein quality control, protein clearance, and intracellular trafficking to various cell-cell recognition events that include cell adhesion, receptor activation, self/non-self-recognition, and host-pathogen interaction^1, 2^. However, much of our knowledge of human glycans is confined to those found on blood cells, free in plasma, and attached to antibodies^3^. We still know very little about cell-specific glycosylation differences in human tissues, including within the lung.

Current evidence suggests human lungs have a very complex glycome, with over 500 *N-*glycan structures regulating tissue development and interactions with inhaled pathogens^3^. Based on their structure, *N*-glycans are classified as pauci-mannose, high-mannose, hybrid, and complex *N*-glycans, which can be additionally decorated with various specialized monosaccharides (i.e., fucose, galactose, and sialic acid) and have distinct branching patterns that will impact their function and interaction with the environment^4^. More specifically, complex *N*-glycans can have bisecting and tetra-antennary organization, polylactosamine and sialic acid decorations, core and antenna fucosylation, or combinations of these^5^. However, the glycan structures involved in cell and tissue physiological and pathological processes remain elusive. This is partly due to insufficient information on their localization, cell type specificity, and protein carriers.

Matrix-assisted laser desorption/ionization mass spectrometry imaging (MALDI-MSI) has become a prime technique for revealing protein *N*-glycan composition and localization within biological tissues^6^. In this approach, *N*-glycans are enzymatically released from their glycoproteins in an in-situ fashion, after which a standard MALDI workflow can be used, where a UV laser is used to ablate and ionize *N*-glycans from the tissues in a spatially defined fashion. Ionized *N*-glycans are then introduced into a mass spectrometer to measure their molecular weight, which indicates their chemical formula and putative identification. This results in the ability to measure and map the relative abundance of these *N*-glycans across tissues in an untargeted, highly multiplexed fashion. Typical applications achieve lateral resolution of tens of microns or less, where approaching cellular resolution in human tissues is possible but not quite there^7^. Compared to lectin and other glycan-binding tissue staining protocols, which can only partially characterize glycan composition (i.e., specific epitope), MALDI-MSI is broad, untargeted, measures an exact mass of the *N*-glycans, and, in combination with specialized databases^8^, reveals *N*-glycans’ composition (e.g., type and number of monosaccharides) in a particular area of the tissue, which is ideal for biomarker discovery and mechanistic studies^9^. For example, we recently discovered that specific *N-*glycans are glomerulosclerosis biomarkers in diabetic kidney disease (DKD) kidneys using MALDI-MSI^7^.

Compatibility of MALDI-MSI-based spatial *N-*glycomics with other imaging modalities affords going beyond just the creation of distributional maps of *N*-glycans to put findings into biological, physiological, or pathological context. The most readily applied approach is performing histochemical (HC) and immunohistochemical (IHC) staining after performing spatial *N*-glycomics, which, for example, revealed *N*-glycan signatures of immune cell populations in lung tissue after COVID-19 infection^10^ and alterations in pulmonary *N*-glycans following irradiation^11^. Further, sequential MALDI-MSI of *N*-glycans and tryptic peptides from the same tissue section enabled the assigning of colocalized protein and *N*-glycan identities that can point to interwoven biochemical processes and can be used as the path toward revealing *N*-glycan protein carriers^12, 13^.

Tissues, such as those comprised in the lung, often have complex architectures and cellular organization. They are composed of various cell types in close association, where multiplexed antibody imaging technologies have become highly desired for single-cell level characterization. One emerging technique that provides a deep view into the single-cell spatial relationships and disease progression is co-detection by indexing (CODEX)^14^, which relies on DNA-conjugated antibodies and the cyclic addition and removal of complementary fluorescently labeled DNA reporters. Compared to traditional IHC, which typically assesses only a limited number of protein markers (two to four) in a tissue section, CODEX can visualize 30 or more markers *in situ*, refining spatial cell type identification^14^.

Herein, we took advantage of the cellular resolution the previously designed CODEX v.2 immunohistochemical imaging panel provides^15-17^, and combined it with MALDI-MSI-based spatial *N*-glycomics on the same human lung tissue section. One limitation in MALDI-MS imaging of *N*-glycans is that signal from one pixel (typically, 20 µm x 20 µm to 100 µm x 100 µm) can often contain content of several neighboring cells, either due to the size of laser ablation spot or unavoidable small delocalization of the *N*-glycans during the sample preparation^18^. While the latest instruments could enable better resolution (down 0.6 μm x 0.6 μm for the transmission mode MALDI-2 source^19^), their application for *N*-glycan imaging was not demonstrated to go beyond 50 µm x 50 µm pixel size^20^. In addition, specific pulmonary cells can be extremely thin, such as the alveolar type 1 cells estimated to be 0.1 µm thick to enable the passive diffusion of oxygen^21, 22^. All these factors limit *N*-glycan discovery studies to tissue functional units rather than specific cells. We previously showed how orthogonal omics measurements, like single-nuclei RNA sequencing from the same tissue, can help determine the cellular origin of *N*-glycan aberrations observed through MALDI-MSI^7^. Here, from the exact same tissue sections, our approach generated protein *N*-glycan profiles of regions comprising many structures and cell types in the normal human lung, and CODEX helped us determine the potential tissue or cellular origin of these protein post-translational modifications. Using our newly developed algorithm for *N*-glycan classification based on their composition, we revealed the chemical properties of *N*-glycans enriched in the specific cell types of a human lung, generating hypotheses for the potential functional role of the specific carbohydrate moieties.

## MATERIAL AND METHODS

### Lung Tissue Source

Transplant-quality donor lung that could not be matched to an organ recipient was obtained through the United Network of Organ Sharing via the National Disease Research Interchange (NDRI) and the International Institute for Advancement of Medicine (IIAM). The organ recovery, storage during transport and processing into the BioRepository for INvestigation of the Lung (BRINDL, https://brindl.urmc.rochester.edu) were performed as described previously^23^ and available at protocol.io^24^. All lung samples were reviewed by pathologists for quality assessment before enrolling in this study. The University of Rochester IRB approved and oversees this study (IRB approval# RSRB00047606).

### Mass spectrometry imaging of *N*-glycans

Step-by-step details of the method can be found in protocols.io^25^. Briefly, a FFPE block of human lung tissue were prepared using the protocol^26^, sectioned at 5 µm thickness and mounted on indium tin oxide (ITO)-coated glass slides. Slides were heated, dewaxed by xylene washes, and rehydrated in serial ethanol (EtOH)/water (v/v) washings and then subjected to antigen retrieval in boiling citraconic buffer followed by PNGase F (N-Zyme Scientifics, 100 µg/mL) spraying using a M5 Sprayer (HTX Technologies), and sample incubation in a relative humidity of 89% for 2 h at 37 °C, as described previously^18^.

After incubation, α-cyano-4-hydroxycinnamic acid (CHCA, Sigma-Aldrich)– 7 mg/mL (50% ACN and 0.1% TFA in water (v/v))– was sprayed over the tissue sections using the M5 Sprayer, as described previously^7^. MALDI-MSI experiments were performed using a scimaX 7 Tesla Magnetic Resonance Mass Spectrometer (MRMS; Bruker Daltonics) equipped with a dual ESI/MALDI ion source and a Smart-beam II Nd:YAG (355 nm) laser. The instrument was operated in 1 w, positive ion mode over an m/z range of 1,000–5,000 with an estimated resolving power of 120,000 at m/z 400. The target plate stepping distance (lateral resolution) was 50 μm. The ion m/z 1809.6393 ([M+Na]^+^ of Hex5 dHex1 HexNAc4) was used as a lock mass for on-line calibration. Imaging data were acquired using FlexImaging (v 4.1, Bruker Daltonics).

### Creating Glycan Mining and Ontology (N-Glycan MiniOn) to automate the dissection of the *N*-Glycan composition

To streamline the classification of the *N*-glycans in the different classes, we developed an R package named Glycan Mining and Ontology (N-Glycan MiniOn) to automate the dissection of the *N*-Glycan composition. This open-source package is available on GitHub (https://github.com/GeremyClair/NglycanMiniOn/). This package is similar to what we previously developed to generate Lipid Ontologies from the lipid names (Lipid MiniOn)^27^. In N-Glycan MiniOn, the function ‘NGlycan_miner()’ parses the names of *N*-Glycans to enable their attribution to different classes. First, it identifies the nature and the exact number of monosaccharides composing the *N*-glycans. Next, using the rules described in **Figure 1G**, it identifies the classes each *N*-glycan belongs to. Once this information is parsed, a second function, ‘Nglycan_ontologies()’, generates a list of ontology terms associated with each *N*-Glycan enabling to perform enrichment analyses using popular methods such as Fisher’s exact test, DAVID’s modified Fisher’s exact test (EASE score)^28^, binomial test, or Kolmogorov–Smirnov test for ranked-based enrichment analyses. It also enables the generation of Figures depicting the frequency of *N*-Glycans in different clusters. A shiny app was developed to streamline the analysis of *N*-glycan structures for non-coding researchers (https://github.com/GeremyClair/NglycanMiniOn_shiny).

**Figure 1.**
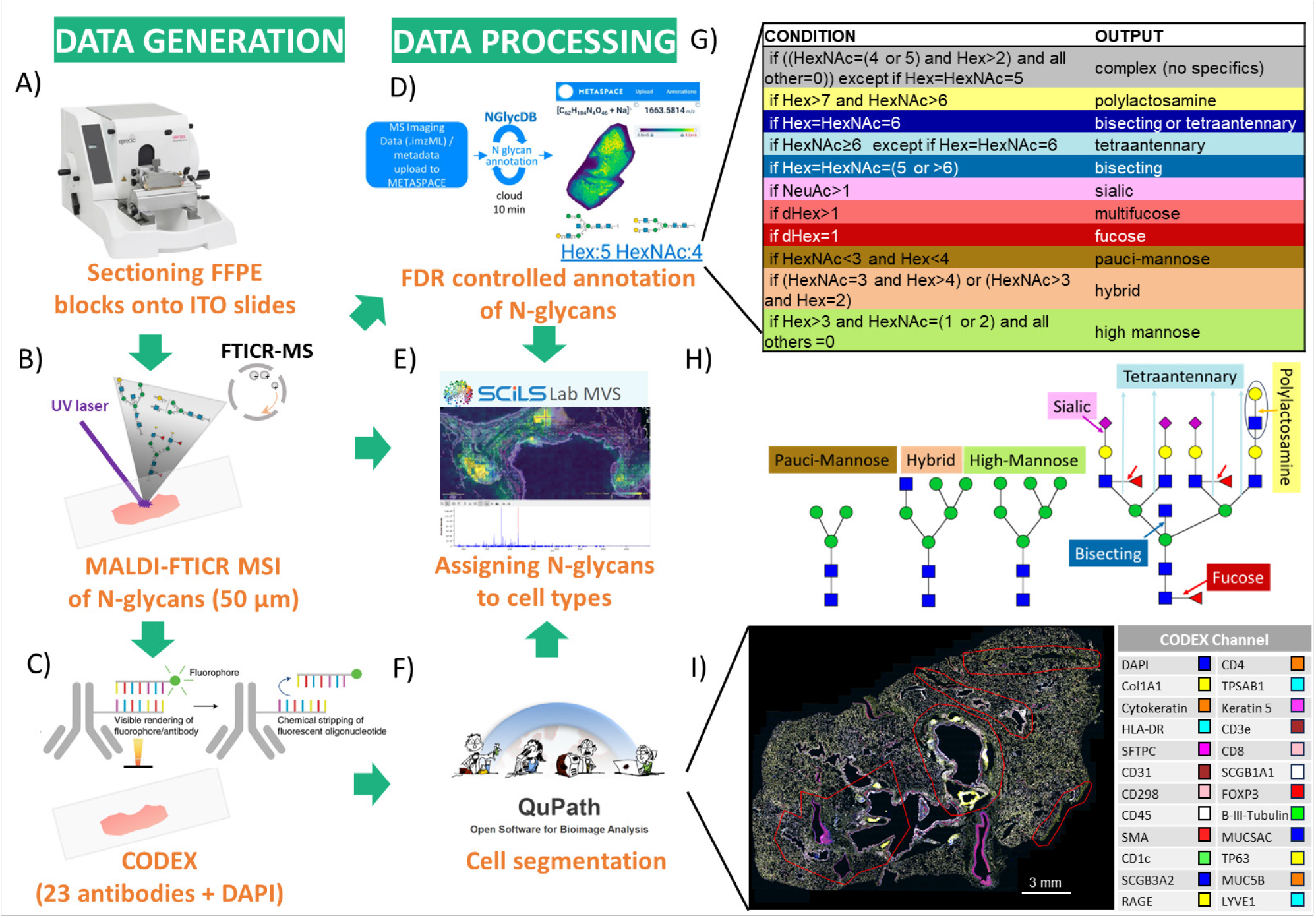
The workflow for integration of MALDI-MSI-based spatial *N*-glycomics and CODEX analysis of the same lung tissue section. (A) The FFPE block of the lung tissue was sectioned, and the same section was used for both (B) *N*-glycan MALDI-MSI and (C) CODEX assays. (D) METASPACE was used for *N*-glycan compositional annotation, and E) SCiLS for integration with (F) CODEX images visualized and segmented using QuPath. (G) Based on their chemical similarities, a classification algorithm for converting *N*-glycan composition to *N*-glycan class was performed to group *N*-glycans. (H) Symbol nomenclature for *N*-glycans highlighting specific structural features. (I) CODEX image, with 24 channels active, generated in QuPath and exported into SCILS for overlaying with *N*-glycan MALDI MSI data. Closed curved red lines in the tissue outline regions where MALDI-MSI was performed.

### CODEX immunohistochemical imaging

Post-MALDI imaging CODEX IHC protocol is available in protocols.io^29^. Briefly, the MALDI-matrix was removed from lung sections through 2 × 2min incubation in 50% ACN and rehydrated via a decreasing ethanol series prior to High pH Heat-Induced Epitope Retrieval (HIER). After cooling, lung tissue was washed with hydration buffer (Akoya Biosciences) and then incubated for 20-30 min in staining buffer (Akoya). Sections were then covered with 200 µL of antibody buffer composed of staining buffer supplemented with recommended concentrations of N, G, J, & S blockers (Akoya Biosciences) and the specified dilutions of up to 34 antibody-barcode conjugates followed by incubation in a humidified chamber at room temperature for 3 h. Sections were then washed with staining buffer followed by fixation in 1.6% paraformaldehyde for 10 min After a series of rinsing in PBS, samples were placed in a storage buffer (Akoya Biosciences) and photobleached by illumination with a 200 mA, 15 watts, 1600 lumens bulb overnight at 4 °C. Image acquisition was performed using the Phenocycler-Fusion 1.0 platform utilizing the 20X (0.5 µm/pixel) objective and the Fusion 1.0.8 software according to manufacturer recommendations, as described^29^.

### Integration of MALDI-MSI and CODEX on the same tissue section

MALDI-MS imaging data files were imported into the SCiLS (Bruker Daltonics) software and exported to imzML. The resulting .imzML and .ibd files were submitted to METASPACE for data processing and *N*-glycan annotation, using the NGlycDB V1 as the database^8^. METASPACE was used for data visualization, where the “Show representative spatial patterns for dataset” tool was used to select distinct spatial patterns. A m/z list of detected *N*-glycans, their annotations at a compositional level, and visualization of their spatial distribution were created using the METASPACE annotation platform, providing information on possible *N*-glycan isomeric structures for each ion image. A m/z list of annotated *N*-glycans was imported back into the SCiLS for integration with CODEX images. CODEX data were opened in QuPath bioimage analysis software and exported as rendered RGB images using available channels^30^. Cell segmentation was performed utilizing the StarDist extension^31, 32^ in QuPath. CODEX images were imported into SCiLS and overlayed with MALDI-MSI ion images created using an annotated m/z list from METASPACE.

## RESULTS AND DISCUSSION

The general concept of our workflow is illustrated in **Figure 1**, where we first performed MALDI-MS imaging of *N*-glycans and then CODEX on the same tissue section. We utilized the METASPACE cloud-based platform for automated molecular annotations and visualization of *N*-glycan data. Our *N*-glycan classification algorithm grouped *N*-glycan annotations based on their chemical similarities. QuPath was used for visualization and segmentation of the CODEX data. Finally, SCiLS was used for the *N*-glycan-CODEX imaging integration.

As a proof-of-concept of our method, we used our optimized *N*-glycan MALDI-MS imaging protocol^18^, which minimizes diffusion (i.e., delocalization of molecules from their endogenous locations) of the released *N*-glycans from their parent glycoprotein, and detected more than 150 *N*-glycans with various spatial patterns associated with different anatomical regions in an FFPE section of the left upper lobe of a human lung. Insight into the compositions, tentative structures, and localization of *N*-glycans can be visualized using the METASPACE platform, where *N*-glycan images are registered and overlayed on a high-resolution microscopy image: https://metaspace2020.eu/project/velickovic-2024_MALDI_CODEX. Manual inspection of data and using the “Show representative spatial pattern for datasets” tool in METASPACE immediately revealed several characteristic spatial patterns of *N*-glycan ion images that align with different anatomical regions in the lung, including adventitial regions of airways and blood vessels, submucosal glands, cartilaginous shields, the smooth muscle of pulmonary artery, and alveolar parenchyma. This spatial heterogeneity and diversity of protein *N*-glycans suggests that these post-translational modifications might play distinct roles in these respective lung tissue functional units.

We then aimed to gain additional insight into the potential structure-function relationship of these protein *N*-glycans. First, using the co-localization analysis, we created a list of *N*-glycans (i.e., their *m/z* and composition) that belong to each spatial structural cell pattern (specifically, anatomical region of the lung) (**Supporting Table S1**). Second, to reveal common *N*-glycan chemical moieties enriched in distinct anatomical/functional regions, we constructed a classification rule (**Figure 1**) that converts the *N*-glycan composition (e.g., number and type of monosaccharides, such as Hex:5 HexNAc:4, **Figure 1D**) to the *N*-glycan structural characteristics (e.g., bisecting, fucosylated, etc.). Thanks to the canonical *N*-glycan structural organization, *N*-glycan composition often reveals much about *N*-glycan structure despite multiple structural isomers that can be associated with the given composition. Herein, we used *N*-glycan compositions and their available structures in NGlycDB within METASPACE^8^ to generate a general classification rule that, based on the number and identity of monosaccharides, defined *N*-glycan as pauci-mannose, hybrid, high-mannose, fucosylated, multi-fucosylated, sialylated, bisecting, tetra-antennary, polylactosamine, or complex without any of these specific moieties. Note that our algorithm does not resolve isomeric structures, but it reveals structural features of protein *N*-glycans that are important to their function, biosynthesis, and antigen properties. One ambiguity occurs with *N*-glycans that, in their composition, contain six hexoses and six *N*-acetylhexosamines. Such *N*-glycans can be ascribed to either bisecting or tetra-antennary structures, and without orthogonal studies, it is impossible to assign correct structural organization. In many cases, *N*-glycan composition implies that its structure contains multiple decorations, where, for example, *N*-glycan Hex:8 HexNAc:8 dHex:2 is classified as a complex bisecting-multifucose-polylactosamine-tetra antennary *N*-glycan.

Using our classification and N-Glycan MiniOn package we developed, we observed that glycoproteins with high mannose glycans are exclusively present in the adventitial region of airways and blood vessels, whereas glycoproteins with complex fucosylated and multifucosylated *N*-glycans are the most numerous structures in the lung. Moreover, some *N*-glycan classes are widespread and colocalize with multiple anatomical regions, including (i) hybrid, complex sialic, and complex multi-fucosylated *N*-glycans that colocalize with the airways and blood vessels adventitial regions, submucosal glands, and cartilage, (ii) complex muti-fucose-tetra antennary structures that colocalize with submucosal glands, cartilage and parenchyma, and (iii) complex fucose-sialic structures that colocalize with airways and blood vessels adventitial regions, submucosal glands, and smooth muscle layers (**Figure 2A**). Other classes of *N*-glycans are however highly specified for certain regions in the lung. For example, glycoproteins with complex fucose-sialic-polylactosamine-tetra-antennary *N-*glycan decorations are highly abundant in smooth muscle cells, while protein *N*-glycans overrepresented in parenchyma include bisecting tetra-antennary structures with and without fucose and/or polylactosamine decorations. Notably, some of these classes are composed of only one glycan structure (**Figure 2A**).

**Figure 2.**
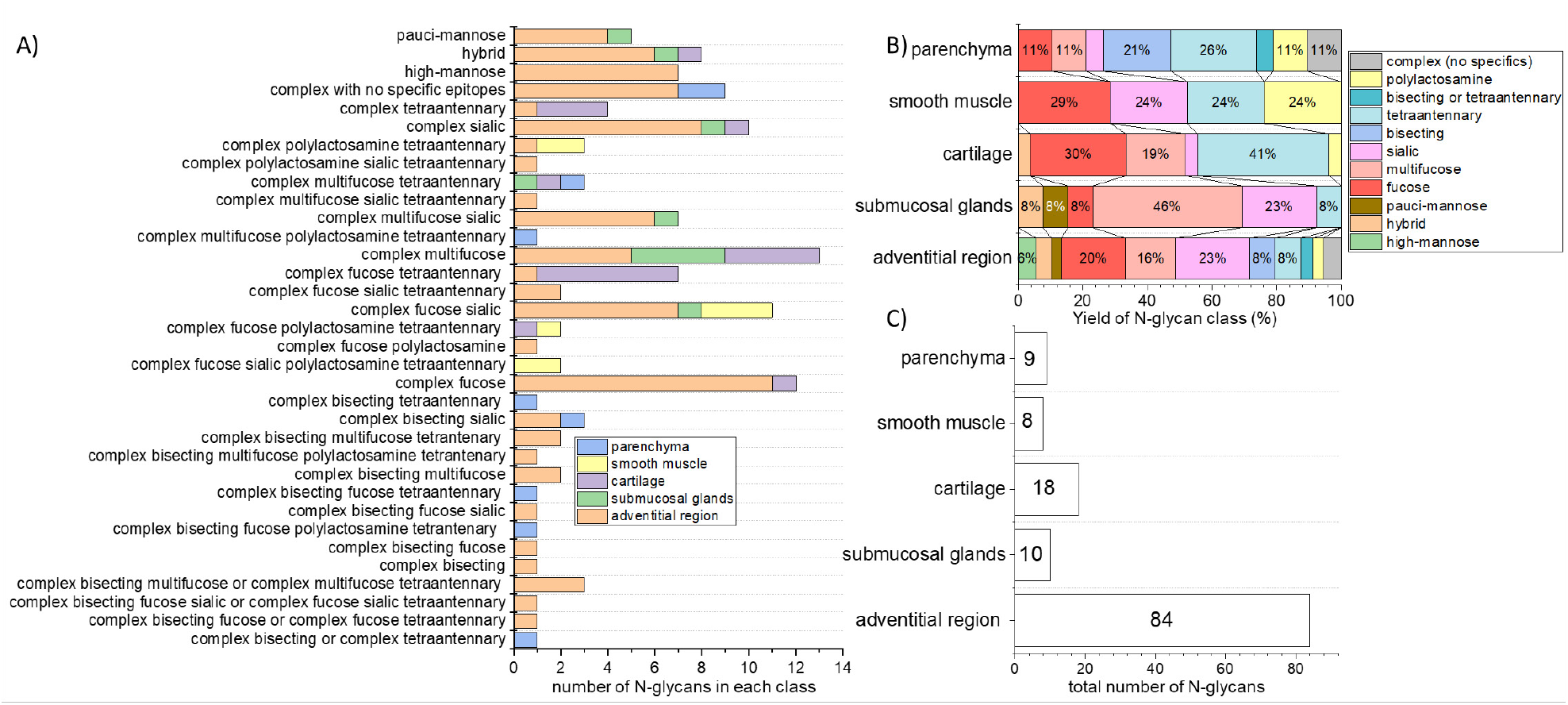
The diversity and abundance of *N*-glycan structures in human lung tissue regions were revealed through MALDI-MSI analysis and the *N*-glycan classification rule. (A) The number of *N*-glycans in each *N*-glycan class detected in each lung section anatomical region. (B) The percentile of each *N*-glycan class in each lung section anatomical region. This was calculated per the total number of *N*-glycans detected in the respective anatomical region. (C) The total number of *N*-glycans in the lung anatomical regions.

Upon investigation of the protein *N*-glycan profile of individual regions (**Figure 2B)**, our data shows that the adventitial region of airways and blood vessels contain the highest diversity of the *N*-glycans, with fucosylated, multi-fucosylated, and sialic acid *N*-glycans representing two-thirds of all such structures. This high diversity of *N*-glycans within the adventitial region is unsurprising, given that we detected the highest number of *N*-glycan structures colocalizing with this anatomical region (**Figure 2C**). On the contrary, smooth muscle cells have only eight highly specialized *N*-glycans that contain fucose, sialic acid, tetraantennary, and polylactosamine structures (or their combinations; **Figure 2C**). Finally, multi-fucosylated *N*-glycans are enriched in the submucosal glands, while cartilage dominates in tetra-antennary structures (**Figure 2B**).

Overlaying MALDI-MS ion images of *N*-glycan distributions with CODEX images acquired from the same section allowed us to decipher the cellular origin of these post-translational modifications in more detail. This was especially true for *N*-glycans surrounding airways, where multiple cell types occur (**Figure 3**). The CODEX image in **Figure 3A** shows protein markers around the bronchus, while the magenta pixels in **Figures 3B-D** show the localization of selected *N*-glycans in the same region. We can observe that multi-fucosylated *N*-glycan (*m/z* 2612.9452) originates from epithelial cells rather than immune cells in submucosal glands (high co-localization of *N*-glycan signal with Pan-cytokeratin marker compared to low co-localization with CD45 marker, **Figure 3B**), while tetra antennary *N*-glycan (*m/z* 1745.6345) originates from cartilage and perichondral fibroblasts where Col1A1 marker is highly expressed (**Figure 3C**). Note, the very bright Col1A1 islands are the result of the MALDI imaging workflow that led to partial detaching, folding, and condensation of cartilaginous plaques (**Supporting Figure S1**), and that in the Col1A1 signal, we can also visualize ablation marks from the MALDI laser. Further, high-mannose *N*-glycans (m/z *1581*.*5283*) surrounding the bronchus (**Figure 3D**) are associated with collagenous structures rather than the adjacent epithelium and smooth muscle cells, as those *N*-glycans were not observed in the vicinity of cells expressing pan-cytokeratin and SMA, respectively.

**Figure 3.**
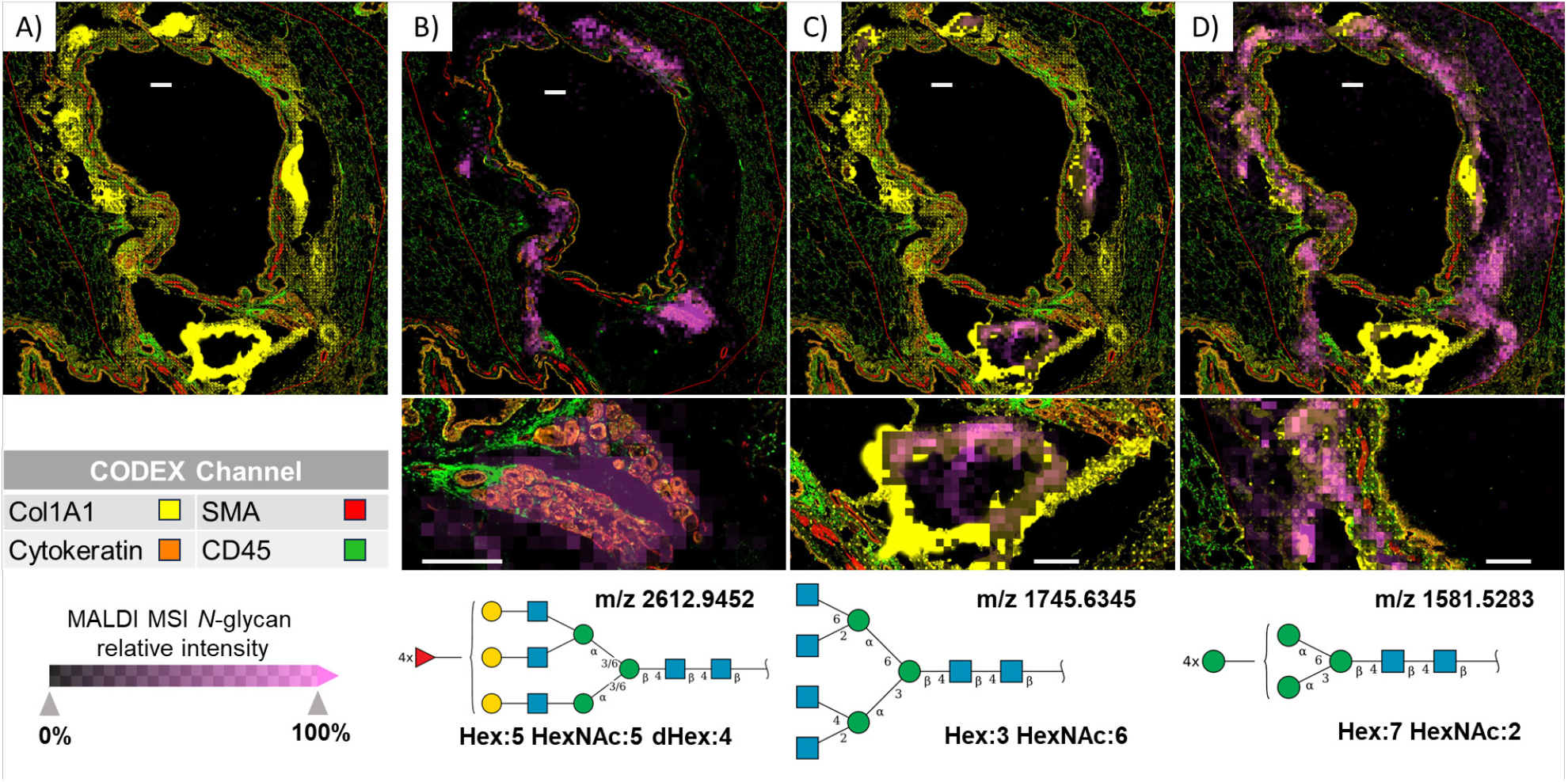
Combining MALDI-MSI and CODEX revealed *N*-glycans and the cell types they are expressed in around the bronchus. A) The CODEX image and CODEX channel color legend B) MALDI-MS image (magenta pixels) of Hex:5 HexNAc:5 dHex:4 (*m/z* 2612.9452) overlaid on CODEX image shows the high abundance of this *N*-glycan in submucosal glands, where Pan-Cytokeratin rather than CD45 is highly expressed. SNFG structure of this complex multifucose *N*-glycan is shown. C) Overlayed MALDI-MS image of Hex:3 HexNAc:6 (*m/z* 1745.6345) over CODEX images shows the highest abundance of this *N*-glycan in cartilage, where Col1A1 is highly abundant. SNFG structure of this complex tetra antennary *N*-glycan is shown. Overlayed MALDI-MS image of Hex:7 HexNAc:2 (*m/z* 1581.5283) over CODEX images shows the high abundance of this *N*-glycan in multiple cell types around the bronchus. SNFG structure of this high-mannose *N*-glycan is shown. White scale bars represent 300 µm.

Inspection of the regions around the pulmonary artery and surrounding parenchyma allowed us to visualize the cellular origin of *N*-glycans with other spatial patterns, **Figure 4**. For example, **Figure 4A-C** reveals that MALDI-MS image pixels of complex fucosylated-tetra antennary-polylactosamine *N*-glycans (*m/z* 3270.1932) are spread over circular regions containing SMA surrounding the airways and arteries, and hence likely smooth muscle structures, but also over epithelium cells distinguished by high expression of pan-cytokeratin. A Hex:6 HexNAc:6 *N*-glycan that, based on our algorithm, can be ascribed to either as a tetra antennary or bisecting *N*-glycan (**Figure 4G**), is highly abundant in the alveolar parenchyma (**Figure 4E-F**), which is characterized by high expression of RAGE and SFTPC markers. However, the spatial resolution of our MALDI-MSI does not allow us to exclude the possibility that *N*-glycans with this spatial pattern are also present in the pulmonary surfactant, in the cells of small airways, or in the cells of the blood vessels closely associated with surrounding parenchyma (**Figure 4E**).

**Figure 4.**
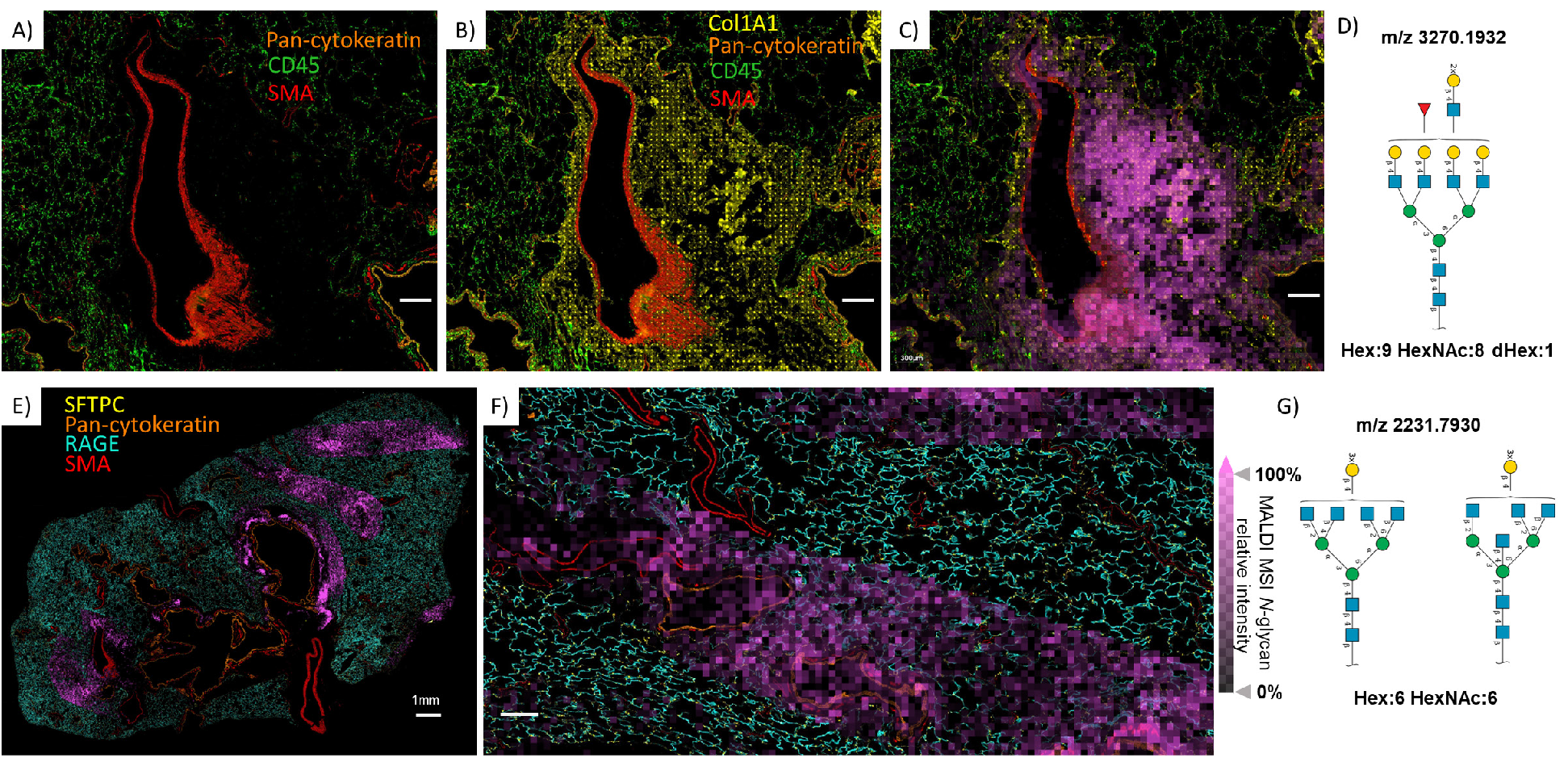
MALDI-MSI *N*-glycans ion images overlaid with CODEX imaging demonstrate region-specific detection of certain *N*-glycans. A) CODEX image performed after the spatial *N*-glycomics analysis shows the location of pan-cytokeratin, CD45, and SMA. B) The same CODEX image as in A with the addition of Col1A1 demonstrates areas of extracellular matrix (ECM) surrounding the pulmonary artery. C) The localization and abundance of *N*-glycan Hex:9 HexNAc:8 dHex:1 in the collagen-rich ECM regions (magenta pixels). D) SNFG structure of this complex-fucosylated-polylactosamine-tetra antennary *N*-glycan. E) and F) illustrate Hex:6 HexNAc:6 post-translational modifications in parenchymal alveolar regions, notably not in the region containing *N*-glycan demonstrated in panel C. G) The SNFG structure of this complex *N*-glycans can be ascribed to as either a tetra antennary or bisecting structure. White scale bars represent 300 µm unless otherwise indicated (Panel E).

In conclusion, we demonstrated the benefits of using CODEX imaging to better interpret the biological context of protein *N*-glycosylation across lung tissue using MALDI-MSI. Specifically, our workflow enables more accurate ascribing *N*-glycans to their cellular and functional tissue unit origin. To fully utilize this approach, we also developed an algorithm that groups *N*-glycans into functional classes based on their compositional information, which allowed us to draw conclusions about the chemistry of *N*-glycans dominant on proteins in the specific functional units of the organ. One drawback of our multimodal approach is that it requires tissue-partially destructive MALDI-MSI to be performed before non-destructive CODEX analysis. This results in visible alterations to the tissue where laser ablated the sample, especially in collagenous structures. Nevertheless, our efforts to perform MALDI-MSI after CODEX suffered a significant drop in *N*-glycan MALDI-MSI sensitivity, so, in our opinion, post-MALDI CODEX is a preferable order. An important note is that correlative *N*-glycan MALDI-MSI and CODEX imaging do not necessarily imply that detected *N*-glycans originate from protein markers, although that is not excluded. In some cases, the MALDI-MSI pixel contains more than one layer of cells identified by CODEX. Therefore, further advancement in spatial resolution of *N*-glycan MALDI-MSI and in developing workflows that enable correlation between detected *N*-glycans, the proteins they modify, and the enzymes involved in their synthesis will unlock the full potential of this novel multimodal imaging approach.

## Notes

**GRANT AND FUNDING INFORMATION:** This work was supported by the National Heart, Lung, and Blood Institute (NHLBI) Molecular Atlas of Lung Development Program (LungMAP) grants U01HL148861 (to G.S.P.) and U01HL148860 (to G.C.), and the Human BioMolecular Atlas Program (HubMAP) grant U54HL165443 (to G.S.P., C.A., and G.C.). Part of this work was performed in the Environmental Molecular Sciences Laboratory (EMSL) from Pacific Northwest National Laboratory, a DOE Office of Science User Facility sponsored by the Office of Biological and Environmental Research and operated under Contract No. DE-AC05-76RL01830.

https://github.com/GeremyClair/NglycanMiniOn/

https://github.com/GeremyClair/NglycanMiniOn_shiny

https://metaspace2020.eu/project/velickovic-2024_MALDI_CODEX?tab=datasets

### Competing Interest Statement

The authors have declared no competing interest.

