## Supporting files for "Integrating *N*-glycan and CODEX imaging reveal cell-specific protein glycosylation in healthy human lung"

**Table of Contents:**

**Figure S1.** Representative spatial patterns of *N*-glycans from METASPACE

**Figure S2.** MALDI-MSI *N*-glycans protocol followed by CODEX highly multiplexed immunofluorescence.

**Tables S1.** List of N-glycans that belong to each spatial pattern observed in METASPACE

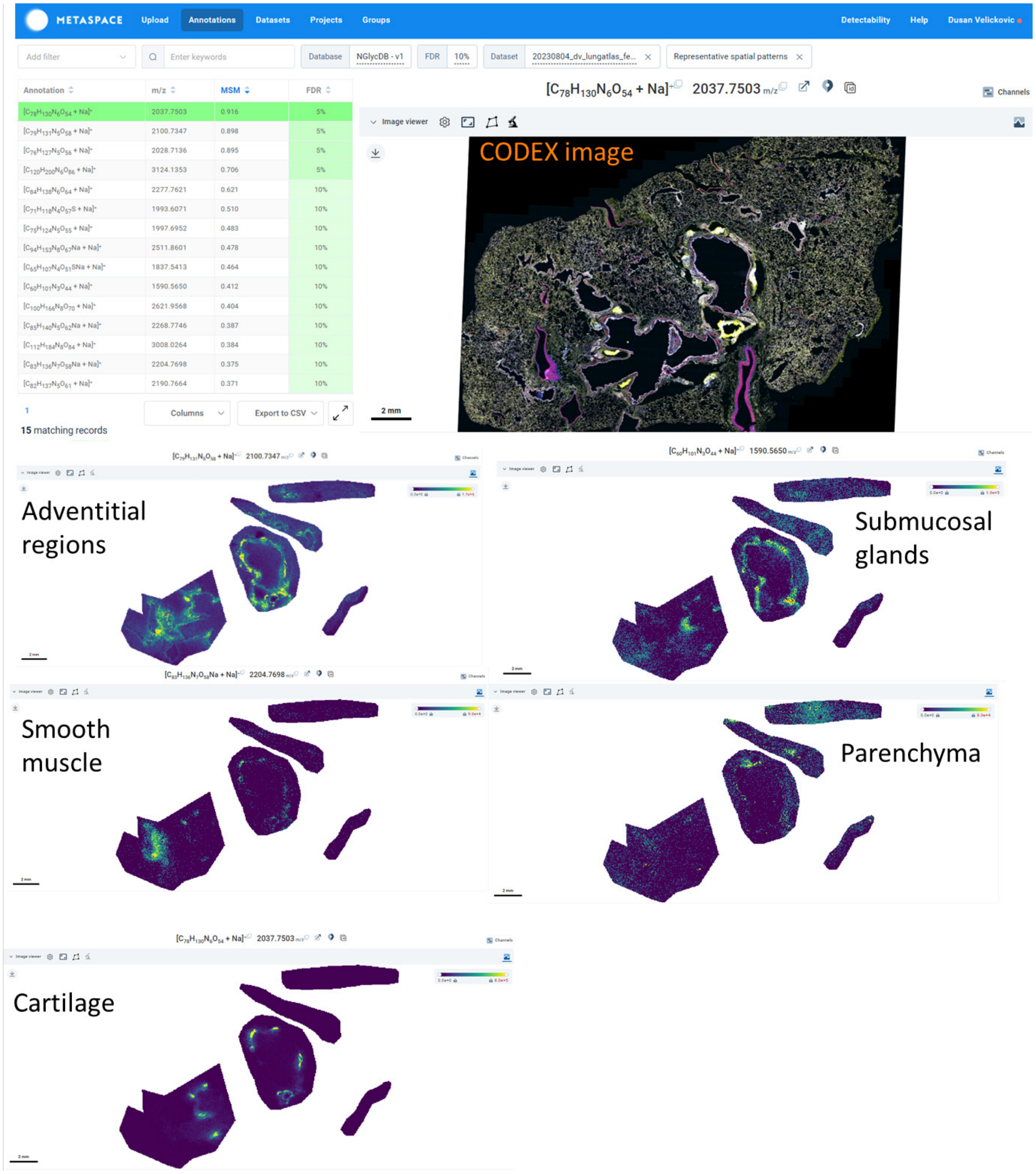


**Figure S1.** Representative spatial patterns of *N*-glycans from METASPACE. 15 spatial patterns were detected (https://metaspace2020.eu/annotations?db_id=353&ds=2023-08-10_00h34m44s&locs=1), with the five most distinct displayed. A list of glycans that co-localize with those features is presented in **Supporting Table S1**.

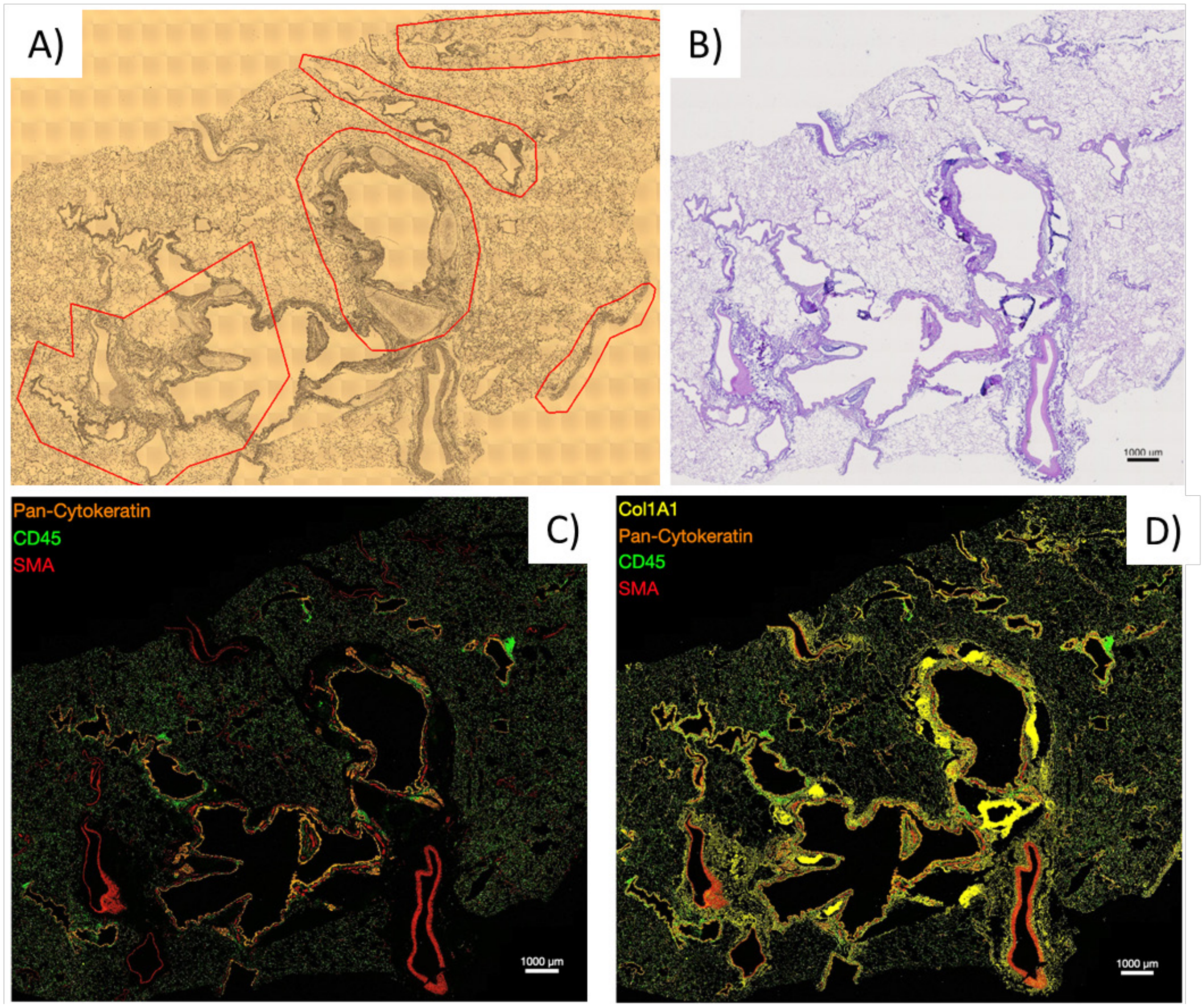


**Figure S2.** MALDI-MSI *N*-glycans protocol followed by CODEX highly multiplexed immunofluorescence. A) Scout image of lung section with regions of interest to be imaged for N-glycans (red objects) and intact cartilaginous plates. B) H&E performed after N-glycan and CODEX assays demonstrate loss and condensation of cartilaginous plates. C) CODEX image of tissue section following *N*-glycans assay showing antibodies for C) pan-cytokeratin, CD45, and SMA and D) with the addition of Col1A1. Collagen is most notable around broncho-vascular structures and particularly dense in folded cartilaginous plates.

**Table S1.** List of N-glycans with different spatial patterns observed in METASPACE. All ions were found as [M+Na] or, in the case of sialic acid, N-glycans, as [M-xH+ (x+1)Na] adducts.

| **Adventitial regions** | |
|---|---|
| **m/z** | **N-glycan composition** |
| 892.2905 | Hex:4 HexNAc:1 |
| 917.3221 | Hex:2 HexNAc:2 dHex:1 |
| 933.317 | Hex:3 HexNAc:2 |
| 974.3435 | Hex:2 HexNAc:3 |
| 1079.375 | Hex:3 HexNAC:2 dHex:1 |
| 1095.37 | Hex:4 HexNAc:2 |
| 1120.402 | Hex:2 HexNAc:3 dHex:1 |
| 1136.396 | Hex:3 HexNAc:3 |
| 1257.423 | Hex:5 HexNAc:2 |
| 1282.454 | Hex:3 HexNAc:3 dHex:1 |
| 1298.449 | Hex:4 HexNAc:3 |
| 1339.476 | Hex:3 HexNAc:4 |
| 1419.476 | Hex:6 HexNAc:2 |
| 1444.507 | Hex:4 HexNAc:3 dHex:1 |
| 1460.502 | Hex:5 HexNAc:3 |
| 1485.534 | Hex:3 HexNAc:4 dHex:1 |
| 1501.529 | Hex:4 HexNAc:4 |
| 1542.555 | Hex:3 HexNAc:5 |
| 1581.528 | Hex:7 HexNAc:2 |
| 1605.54 | Hex:4 HexNAc:3 NeuGc:1 |
| 1606.56 | Hex:5 HexNAc:3 dHex:1 |
| 1622.555 | Hex:6 HexNAc:3 |
| 1630.571 | Hex:3 HexNAc:4 NeuAc:1 |
| 1647.587 | Hex:4 HexNAc:4 dHex:1 |
| 1663.581 | Hex:5 HexNAc:4 |
| 1688.613 | Hex:3 HexNAc:5 dHex:1 |
| 1704.608 | Hex:4 HexNAc:5 |
| 1743.581 | Hex:8 HexNAc:2 |
| 1792.624 | Hex:4 HexNAc:4 NeuAc:1 |
| 1793.644 | Hex:4 HexNAc:4 dHex:2 |
| 1809.639 | Hex:5 HexNAc:4 dHex:1 |
| 1850.666 | Hex:4 HexNAc:5 dHex:1 |
| 1866.661 | Hex:5 HexNAc:5 |
| 1905.634 | Hex:9 HexNAc:2 |
| 1938.682 | Hex:4 HexNAc:4 dHex:1 NeuAc:1 |
| 1954.677 | Hex:5 HexNAc:4 NeuAc:1 |

| Adventitial regions (cont.) | |
|---|---|
| **m/z** | **N-glycan composition** |
| 1955.697 | Hex:5 HexNAc:4 dHex:2 |
| 1971.692 | Hex:6 HexNAc:4 dHex:1 |
| 1995.703 | Hex:4 HexNAc:5 NeuAc:1 |
| 2012.719 | Hex:5 HexNAc:5 dHex:1 |
| 2028.714 | Hex:6 HexNAc:5 |
| 2067.687 | Hex:10 HexNAc:2 |
| 2100.735 | Hex:5 HexNac:4 dHex:1 NeuAc:1 |
| 2101.755 | Hex:5 HexNAc:4 dHex:3 |
| 2157.756 | Hex:5 HexNAc:5 NeuAc:1 |
| 2158.777 | Hex:5 HexNAc:5 dHex:2 |
| 2174.772 | Hex:6 HexNAc:5 dHex:1 |
| 2245.772 | Hex:5 HexNAc:4 NeuAc:2 |
| 2303.814 | Hex:5 HexNAc:5 dHex:1 NeuAc:1 |
| 2304.835 | Hex:5 HexNAc:5 dHex:3 |
| 2320.829 | Hex:6 HexNAc:5 dHex:2 |
| 2336.824 | Hex:7 HexNAc:5 dHex:1 |
| 2341.791 | Hex:6 HexNAc:5 NeuAc:1 |
| 2377.851 | Hex:6 HexNAc:6 dHex:1 |
| 2391.83 | Hex:5 HexNAc:4 dHex:1 NeuAc:2 |
| 2392.851 | Hex:5 HexNAc:4 dHex:3 NeuAc:1 |
| 2393.846 | Hex:7 HexNAc:6 |
| 2466.887 | Hex:6 HexNAc:5 dHex:3 |
| 2487.849 | Hex:6 HexNAc:5 dHex:1 NeuAc:1 |
| 2522.888 | Hex:6 HexNAc:6 NeuAc:1 |
| 2523.909 | Hex:6 HexNAc:6 dHex:2 |
| 2537.888 | Hex:5 HexNAc:4 dHex:2 NeuAc:2 |
| 2539.904 | Hex:7 HexNAc:6 dHex:1 |
| 2668.946 | Hex:6 HexNAc:6 dHex:1 NeuAc:1 |
| 2669.967 | Hex:6 HexNAc:6 dHex:3 |
| 2756.962 | Hex:6 HexNAc:5 dHex:1 NeuAc:2 |
| 2757.983 | Hex:6 HexNAc:5 dHex:3 NeuAc:1 |
| 2758.978 | Hex:8 HexNAc:7 |
| 2816.025 | Hex:6 HexNAc:6 dHex:4 |
| 2830.999 | Hex:7 HexNAc:6 dHex:1 NeuAc:1 |
| 2832.02 | Hex:7 HexNAc:6 dHex:3 |
| 2887.025 | Hex:5 HexNAc:5 dHex:3 NeuAc:2 |
| 2889.041 | Hex:7 HexNAc:7 dHex:2 |
| 2903.02 | Hex:6 HexNAc:5 dHex:2 NeuAc:2 |
| 2905.036 | Hex:8 HexNAc:7 dHex:1 |

| Adventitial regions (cont.) | |
|---|---|
| **m/z** | **N-glycan composition** |
| 3035.099 | Hex:7 HexNAc:7 dHex:3 |
| 3050.073 | Hex:8 HexNAc:7 NeuAc:1 |
| 3122.095 | Hex:7 HexNAc:6 dHex:1 NeuAc:2 |
| 3254.173 | Hex:8 HexNAc:8 dHex:2 |
| 3269.173 | Hex:7 HexNAc:6 dHex:4 NeuAc:1 |
| **Submucosal glands** | |
| **m/z** | **N-glycan composition** |
| 1225.433 | Hex:3 HexNAc:2 dHex:2 |
| 1428.512 | Hex:3 HexNAc:3 dHex:2 |
| 1589.545 | Hex:4 HexNAc:3 NeuAc:1 |
| 1590.565 | Hex:4 HexNAc:3 dHex:2 |
| 1768.613 | Hex:6 HexNAc:3 dHex:1 |
| 1776.629 | Hex:3 HexNAc:4 dHex:1 NeuAc:1 |
| 2246.793 | Hex:5 HexNAc:4 dHex:2 NeuAc:1 |
| 2247.813 | Hex:5 HexNAc:4 dHex:4 |
| 2612.945 | Hex:6 HexNAc:5 dHex:4 |
| 2978.077 | Hex:7 HexNAc:6 dHex:4 |
| **Cartilage** | |
| **m/z** | **N-glycan composition** |
| 1323.4808 | Hex:2 HexNAc:4 dHex:1 |
| 1631.5916 | Hex:3 HexNAc:4 dHex:2 |
| 1833.6505 | Hex:3 HexNAc:5 NeuAc:1 |
| 1745.6345 | Hex:3 HexNAc:6 |
| 1891.6924 | Hex:3 HexNAc:6 dHex:1 |
| 2037.7503 | Hex:3 HexNAc:6 dHex:2 |
| 2183.8082 | Hex:3 HexNAc:6 dHex:3 |
| 2297.8511 | Hex:3 HexNAc:8 dHex:1 |
| 1996.7238 | Hex:4 HexNAc:5 dHex:2 |
| 2053.7452 | Hex:4 HexNAc:6 dHex:1 |
| 2110.7667 | Hex:4 HexNAc:7 |
| 2256.8246 | Hex:4 HexNAc:7 dHex:1 |
| 2069.7401 | Hex:5 HexNAc:6 |
| 2215.798 | Hex:5 HexNAc:6 dHex:1 |
| 2361.856 | Hex:5 HexNAc:6 dHex:2 |
| 2621.9568 | Hex:5 HexNAc:8 dHex:1 |
| 2580.9302 | Hex:6 HexNAc:7 dHex:1 |
| 3311.1946 | Hex:8 HexNAc:9 dHex:1 |

| **Smooth muscle** | |
|---|---|
| **m/z** | **N-glycan composition** |
| 1938.6819 | Hex:4 HexNAc:4 dHex:1 NeuAc:1 |
| 2141.7613 | Hex:4 HexNAc:5 dHex:1 NeuAc:1 |
| 2204.7698 | Hex:3 HexNAc:6 dHex:1 NeuAc:1 |
| 3124.1102 | Hex:9 HexNAc:8 |
| 3196.1313 | Hex:8 HexNAc:7 dHex:1 NeuAc:1 |
| 3270.1681 | Hex:9 HexNAc:8 dHex:1 |
| 3583.2454 | Hex:9 HexNAc:8 dHex:1 NeuAc:1 |
| 3635.3003 | Hex:10 HexNAc:9 dHex:1 |
| **Parenchyma** | |
| **m/z** | **N-glycan composition** |
| 1825.6342 | Hex:6 HexNAc:4 |
| 1987.687 | Hex:7 HexNAc:4 |
| 2157.7562 | Hex:5 HexNAc:5 NeuAc:1 |
| 2231.793 | Hex:6 HexNAc:6 |
| 2596.9252 | Hex:7 HexNAc:7 |
| 2685.9616 | Hex:7 HexNAc:6 dHex:2 |
| 2742.9831 | Hex:7 HexNAc:7 dHex:1 |
| 3051.0938 | Hex:8 HexNAc:7 dHex:2 |
| 3108.1153 | Hex:8 HexNAc:8 dHex:1 |
